# When can AlphaFold predict the oligomeric states of proteins?

**DOI:** 10.1101/2025.03.10.642518

**Authors:** Yiechang Lin, Ciara Wallis, Ben Corry

## Abstract

Homooligomerisation is a prevalent and important process that many proteins undergo to form the quaternary structures required for biological function. However, determining oligomeric states and structures experimentally remains technically challenging and time-consuming for many proteins. Here, we show that the protein structure prediction tools AlphaFold2-Multimer and AlphaFold3 can be used to quickly and accurately predict oligomeric states and structures for a range of soluble and membrane proteins. Using a benchmark set of 40 proteins, we provide optimal parameters for minimizing computational cost while maintaining accuracy. We further extended this analysis to a large dataset of over 1,000 proteins using AlphaFold2-Multimer and observe comparable overall performance but find that accurate oligomeric state prediction remains challenging for proteins that lack close structural representatives in the AlphaFold training set. Together, our results suggest both the utility and current limitations of AlphaFold-based oligomeric state prediction, highlight cases where multiple physiologically relevant assemblies may be plausible, and provide practical guidance for applying these methods to proteins lacking experimental structural data.

## Introduction

Between 30-50% of all proteins are thought to self-associate into stable complexes consisting of multiple copies of themselves.^1,2^ This oligomerisation process is important for the structural and functional properties of these proteins, as it enables the formation of repetitive structural elements, catalytic interfaces in enzymes and the central pore in many membrane proteins. Establishing the biologically relevant oligomeric state of proteins, and the structures they adopt is thus critical for understanding the way in which they perform their biological roles.

A variety of biochemical and biophysical techniques such as native mass spectrometry, size exclusion chromatography and native gel electrophoresis are available to characterize the oligomeric states of proteins.^3^ However, these experimental techniques can sometimes be expensive, time-consuming and limited by technical constraints. Existing computational tools to characterize protein oligomerisation states include sequence similarity-based methods which predict the oligomerisation state using related proteins with resolved structures.^4^ Once the number of subunits is known, a model of the oligomer can be built using a structural templating technique if a relevant template exists or ab initio docking where this is not possible. However, this method relies on the existence of oligomeric state information from a related protein. More recently developed deep learning tools trained on large datasets of curated subunit information from experimental data predicts oligomeric states with relatively high accuracy but do not provide structural information.^5,6^ Currently, for yet to be characterized proteins, the AlphaFold Protein Structure Database (AFDB) provides predicted monomeric structures but lacks predictions for functionally relevant multimeric states which may provide key insights into their physiological behaviour.^7^

Since its release, AlphaFold2 Multimer has found wide application – including to identify plant-pathogen interactions across two different species,^8^ to predict protein-protein complexes in yeast,^9^ and capture interactions between intrinsically disordered protein regions.^10^ Additionally, recent proteome wide predictions using AlphaFold2 highlighted the high proportion of proteins which form homomers and highlight the potential for structural prediction tools to model protein homooligomerization.^2^ AlphaFold3 provides further improvements in accuracy for the modelling of protein-protein interactions, in addition to the ability to model structures of other biological molecules.^11^ More recently, AlphaFold and AlphaFold based tools have also been applied to predict oligomeric states and model structures of homomeric complexes across different datasets.^12–14^

Here, we investigate whether AlphaFold-Multimer v2.3 (AF2-M) and AlphaFold3 (AF3) can correctly predict both the oligomeric states of proteins and the corresponding multimeric structures. Using a curated benchmark set of 40 proteins, we show that the interface predicted TM (ipTM) score reported by both methods is a reliable indicator of oligomeric state, and that the resulting multimeric structures closely resemble experimentally resolved assemblies. Based on this analysis, we identify prediction strategies that minimise computational cost while maintaining accuracy for proteins with unknown oligomeric states.

We then extend this approach to a dataset of over 1,000 proteins and observe a similar overall level of accuracy in oligomeric state prediction following filtering to exclude low-confidence structural predictions. However, we find that proteins lacking structures or close homologues in the AlphaFold training set are substantially less likely to be predicted in the correct oligomeric state, even when their predicted structures are of high confidence. Finally, we apply this technique to a select set of membrane proteins and show that small differences in ipTM scores between multiple oligomeric states can indicate proteins that may adopt more than one biologically relevant assembly.

## Results

First, we asked whether the ipTM score reported by AF2-M, which measures the accuracy of the predicted relative positions of subunits within a protein-protein complex, can be used to correctly predict known protein oligomeric states.

Using the dataset curated from the UniProt Database by Deng et al. (2024), for their study on oligomeric state prediction using a deep learning approach,^6^ we extracted a sample of 40 proteins with known oligomeric states (Table S1). This sample was selected to contain 8 examples each of proteins known to have oligomeric states of 1, 2, 3, 4 and 5 and contained a mix of soluble and membrane proteins (6 single-pass and 10 multi-pass). For each protein, we used AF2-M to predict their structures as homodimers, homotrimers, homotetramers and homopentamers. We used 3 recycles and predicted 20 structures in each oligomeric state for a total of 80 structures produced per protein.

For the 32 multimeric proteins in our dataset, the distribution of ipTM scores for all structures predicted in the correct oligomeric state (e.g., a known dimeric protein predicted with two subunits) differs considerably from the distribution found for all structures predicted in an incorrect oligomeric state (Fig. 1A). In general, higher ipTM scores are observed (peak at ipTM 0.75-1) for proteins predicted in their annotated oligomeric states compared to an incorrect state (peak at ipTM 0-0.32). While there exists some overlap between the distributions, the presence of distinct differences between these peaks suggests the ipTM score reported by AF2-M can distinguish between correctly and incorrectly predicted oligomeric states. For the 8 known monomeric proteins in our test set, predicting them as homodimers, homotrimers, homotetramers, or homopentamers consistently yielded low ipTM scores (0-0.4). The distribution of their ipTM scores differs substantially to that of the known 32 known multimeric proteins predicted in any oligomeric state (Fig. 1B), suggesting that AF2-M can also distinguish between proteins likely to be monomeric and those forming higher order protein assemblies.

**Figure 1.**
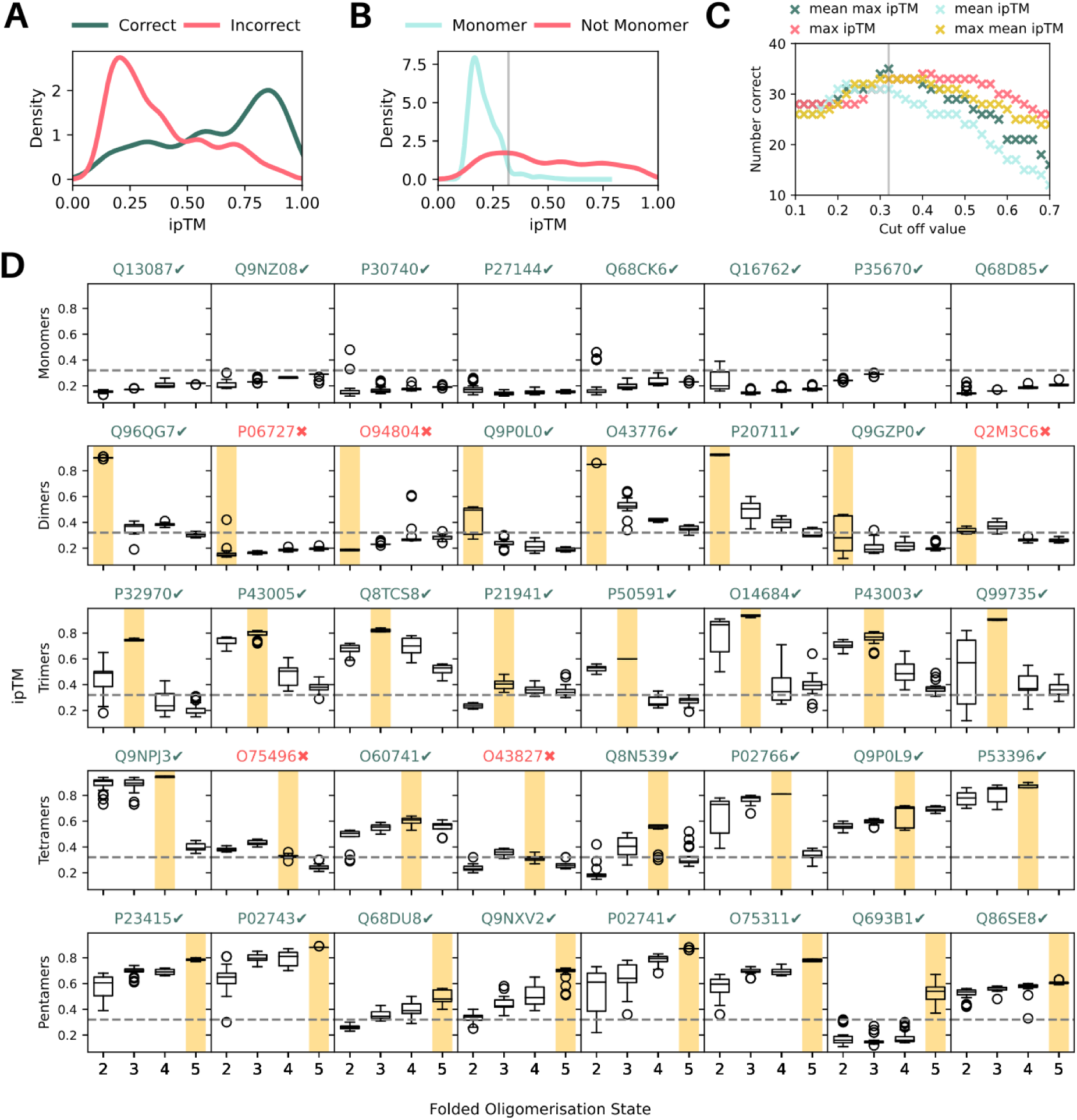
ipTM scores from AlphaFold2 Multimer correctly predict oligomeric states. (A) Probability distribution plot of ipTM scores for 32 known multimeric proteins predicted in the correct (green) and incorrect (pink) oligomeric states. (B) Probability distribution plot of ipTM scores for 8 known monomeric proteins predicted with 2,3,4,5 subunits (blue) and 32 known multimeric proteins predicted with 2,3,4,5 subunits (pink). The grey line indicates the cut-off ipTM value of 0.32. (C) Number of correctly assigned oligomeric states (out of 40) vs cut-off value selected for 4 different methods to distinguish between monomeric and multi-subunit proteins. The grey line indicates the optimal cut-off ipTM value of 0.32. (D) Box plots showing individual ipTM scores for different folded oligomeric states across the test set of 40 proteins with the correct oligomeric state highlighted in yellow. The dashed line indicates the cut-off ipTM value of 0.32. UniProt IDs are noted in green for correct predictions and red for incorrect ones.

These data indicate that we may be able to predict a protein’s most likely oligomeric state by selecting the state which yields the highest mean or max ipTM score (calculated across the 20 structures predicted for each protein) if we can firstly define a cut-off value to assign a protein as monomeric, since monomeric predictions do not have an associated ipTM score. A cut-off value that is too low would lead to incorrect assignment of some monomeric proteins as multimeric and a too high cut-off value could lead to incorrect assignment of some multimeric proteins as monomeric. We optimised the cut-off value across four methods – taking the mean of all ipTM scores for a protein (mean), using the mean of the highest ipTM value for each oligomeric state (mean max), using the maximum of the mean ipTM value for each oligomeric state (max mean) and taking the maximum ipTM score for the protein (max) (Fig. 1C). We found that taking the mean of the maximum ipTM score of each oligomeric state for the protein lead to the highest number of correct states assigned when we optimise this cut-off value to 0.32 and use the max ipTM to assign the oligomeric state if it is not a monomer.

Using this method, we find that 35/40 proteins in our dataset are correctly assigned. Individual ipTM distributions across the 40 proteins highlights trends across the different types of oligomeric states (Fig. 1D). Across monomeric proteins, predictions at all oligomeric states yield low ipTM scores. In this set of proteins, we find that all monomeric, homotrimeric and homopentameric proteins were assigned to the correct oligomeric state. Three homodimers were incorrectly assigned – as a monomer (UniProt ID: P06727), tetramer (UniProt ID: O94804) and trimer (UniProt ID: Q2M3C6). Additionally, two homotetramers were incorrectly assigned as trimers (UniProt IDs: O7549, O43827). Interestingly, all incorrectly assigned proteins had low ipTM scores across all oligomeric states, even those that were not ultimately assigned as monomeric, suggesting that the absolute values of the ipTM scores may be used to gauge confidence in the prediction if a protein is predicted as a homooligomer.

We note that the annotated oligomeric states for each protein typically have smaller ipTM ranges when compared to other oligomeric states (Fig. S1A). Additionally, the average pLDDT (a per-residue measure of local confidence) cannot be used to distinguish between annotated and other oligomeric states, as the monomeric form usually has the highest pLDDT score (Fig. S1B).

Next, we examined whether the structures of the oligomeric assemblies predicted resembled the monomeric structures in the AFDB and experimentally determined structures. Snapshots of the top ranked structural predictions in the correct oligomeric state were produced for each protein after aligning to the monomeric AFDB structure (Fig. 2A).

**Figure 2.**
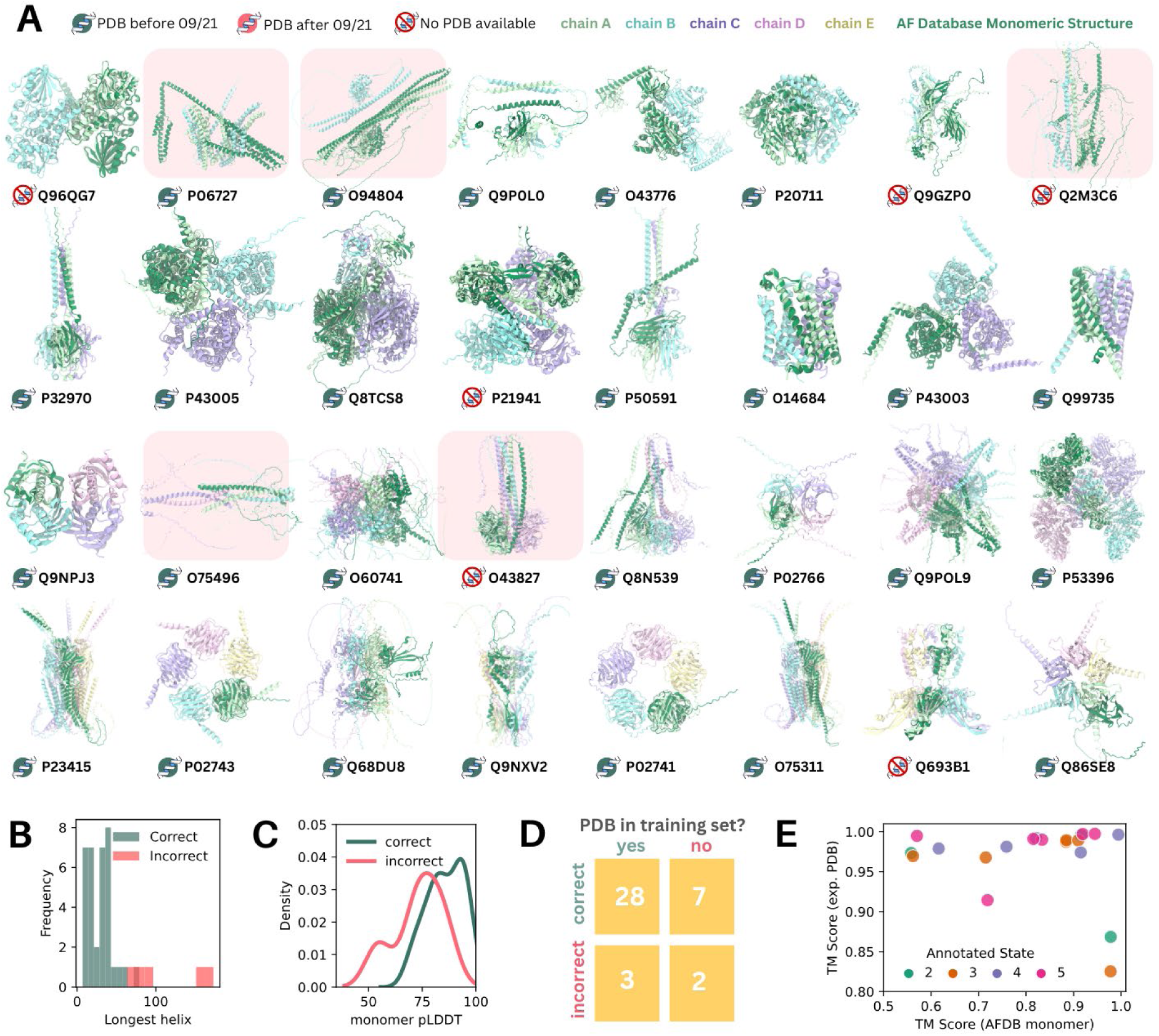
Multimeric structures predicted by AlphaFold2 resemble experimentally determined structures. (A) Snapshots of the top ranked structural predictions in the correct oligomeric state for each multimeric protein (ribbon, coloured by chain), aligned with the AFDB prediction of the protein as a monomer (ribbon, dark green). The presence of an experimentally resolved structure in the PDB or AlphaFold2 training set is indicated with an icon near each UniProt ID. The structures of proteins predicted in an incorrect oligomeric state are highlighted in red. (B) Histogram showing the length of the longest helix in the corresponding AFDB monomer structure for proteins predicted in the correct (green) and in correct (pink) oligomeric states. (C) Probability distribution plot of average monomeric pLDDT values (using the AFDB structure) for proteins predicted with the correct (green) and incorrect (pink) oligomeric states. (D) Two-way table showing the distribution of proteins with oligomeric states predicted correctly and incorrectly based on whether an experimentally resolved structure exists within the AlphaFold2 training set. (E) TM score between the top ranked AlphaFold2 predicted structure and experimentally resolved structure (if available) plotted against the TM score between the top ranked AlphaFold2 predicted structure and the monomeric structure from the AFDB, coloured by annotated oligomerisation state.

Proteins where the oligomeric state was incorrectly predicted by AF2-M all have abnormally long helices in both the monomeric and oligomeric structures (Fig. 2A-B), with the longest helices spanning 64-173 residues. We examined also whether the average monomeric pLDDT, calculated using the structure present in the AFDB indicates whether the correct oligomeric state will be predicted (Fig. 2C). In this sample set, if the pLDDT value was > 90 (14 structures), then the correct oligomeric state was assigned. If the pLDDT value was < 50 (1 structure), then an incorrect oligomeric state was predicted.

The presence of an experimentally resolved structure in the AlphaFold2.3 training set (prior to 30 September 2021) did not have a large impact on whether the correct oligomeric state was predicted in this set of proteins (Fig. 2D). 28/31 of structures present in the training set were correctly predicted compared to 7/9 structures absent from the training set being correctly predicted.

Next, we quantified the similarity between the predicted multimeric structures to both experimentally resolved structures and to AF2 monomeric predictions found in the AFDB in the cases for which the oligomeric state was correctly predicted and an experimentally resolved structure exists (Fig. 2E, Table S2). The TM score between the computationally predicted and experimentally determined structures are generally very high across all oligomerisation states, with most predicted structures having TM scores of greater than 0.95 to experimentally resolved structures. This suggests that predicted structures of higher order oligomers are likely to be accurate. In many cases, the predicted structures of subunits in the multimer showed significant differences to the AF2 predicted monomeric structure (Fig, 2E), suggesting changes to the individual subunit structure imposed by oligomer formation. Visual analysis of these cases (e.g. UniProt ID: P5O591) indicates that this is usually due to changes in the relative orientations of protein domains and highlights a limitation of using predicted monomeric structures from the AFDB.

Taken together, this suggests that AF2-M can predict the oligomeric states and structures of homomeric proteins accurately, independent of whether the protein structure is present in the training set. The oligomeric state prediction is likely to be accurate if the monomeric pLDDT is high and there are no abnormally long alpha helices in the monomeric structure.

To optimise the computational cost associated with making these predictions, which can be considerable for larger proteins, we next assessed how the chosen number of recycles and number of predicted structures influences the accuracy of oligomeric state prediction. We carried out predictions using combinations of 0, 1, 2, 3 recycles and 1, 5, 10, 20 structures per condition using same set of proteins and assessed how these parameters altered the ipTM scores of the structures predicted in the annotated and incorrect oligomeric states.

Increasing the number of structures from 1 to 5, 10 or 20 does not generally increase the mean ipTM score per protein (Fig. 3A). This is true for predictions of proteins in both the annotated (green, top row) and other (red, bottom row) oligomeric states. However, for a few proteins (outlier values in box plots), increases in ipTM by up to 0.2 are observed when 5, 10 or 20 structures are predicted when compared to 1 structure. Increasing the number of recycles to 1, 2 or 3 generally increases the mean ipTM score compared to 0 recycles, particularly for structures predicted in their annotated (green, top) oligomeric states with 5, 10 or 20 structures (Fig. 3B). The mean ipTM score for proteins predicted in incorrect oligomeric states (red, bottom) also generally increases, however, for some proteins, additional recycles decreases the mean ipTM score instead.

**Figure 3.**
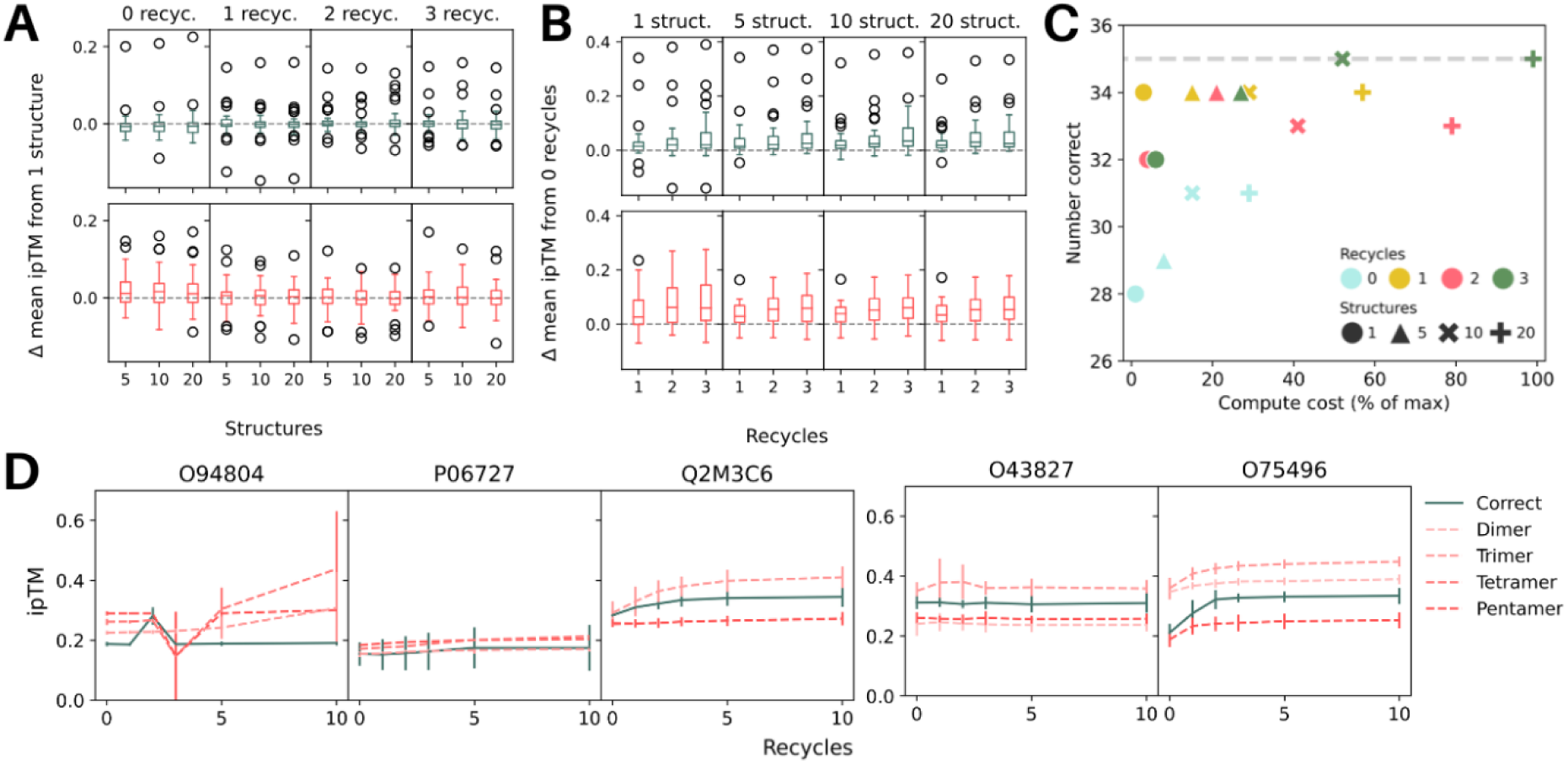
Optimising the computational cost. (A) Distribution of change in mean ipTM from 1 structure and either 0,1,2,3 recycles versus the number of predicted structures for proteins folded in the same oligomeric state as annotated (top, green) and in a different state than annotated (bottom, pink). Each datapoint represents one protein. (B) Distribution of change in mean ipTM from 0 recycles and either 1,5,10 and 20 structures versus the number of recycles predicted for proteins folded in the same oligomeric state as annotated (top, green) and in a different state than annotated (bottom, pink). Each datapoint represents one protein. (C) Number of correct oligomeric state predictions (using monomeric cut off value of 0.32) vs % of max compute cost across 0,1,2,3 recycles and 1,5,10,20 structures predicted. The dashed grey line indicates the highest number of correct predictions (35) across all parameters. (D) ipTM scores of the annotated (green) and incorrect (red, various line styles) states across 0, 1, 2, 3, 5, and 10 recycles for incorrectly assigned dimers (left) and incorrectly assigned tetramers (right).

The comparison between compute cost and number of proteins predicted in the correct oligomeric state suggests that 3 recycles and 10 structures is the most computationally efficient set of parameters, predicting 35/40 proteins in the correct oligomeric state (Fig. 3C). Producing 5 structures with 1,2, or 3 recycles also each yielded 34/40 correct predictions for a lower compute cost. The worst results are obtained using no recycles.

For the five proteins which were predicted incorrectly using 3 recycles and 20 structures, increasing the number of recycles to 5 or 10 does not correct the prediction and ipTM scores for structures predicted in both the incorrect and annotated oligomeric states do not further increase after 5 recycles (Fig. 3D).

While use of the raw ipTM metric was relatively successful for an initial benchmark set of 40 proteins, we next evaluated its performance at predicting the correct oligomeric state with a substantially larger and more challenging dataset. While some proteins in the benchmark set lacked experimentally resolved structures, a similarity search with Foldseek^15^ revealed that highly similar homologous structures were present in the AlphaFold training set for these proteins (Table S1), raising the question of how strongly accurate oligomeric state prediction depends on the presence of close structural homologues in the training data. To explicitly test this, we randomly selected 1,006 proteins from the DeepSub database with annotated homo-oligomeric states, none of which had experimental structures present in the AlphaFold training set, reducing the possibility of template memorisation. Each protein was modelled as a dimer, trimer, tetramer, and pentamer using AF2-M, and oligomeric state prediction was assessed based on relative ipTM values across these assemblies.

Proteins were divided into two classes, “seen-like” and “unseen”, based on the Foldseek determined TM score between the AFDB monomeric structure and experimentally determined structures in the AlphaFold training set. “Seen-like” proteins had at least one homologous structure with a Foldseek TM score > 0.5, whereas “unseen” proteins showed no detectable similarity to any PDB structure (TM score < 0.5), corresponding to proteins for which AlphaFold has no close structural reference.

Compared to the initial benchmark set, the overall proportion of proteins predicted correctly by AF2-Multimer was slightly lower for both seen-like and unseen proteins (80.7% and 66.4%, respectively). However, filtering predictions by pLDDT, using either the monomeric structure or the predicted oligomeric complex, improved accuracy for both groups, with maximum accuracy occuring for structures where the pLDDT of the oligomeric complex was above 90 (88.5% for seen-like proteins, and 82% for unseen proteins) (Fig. 4A, Fig. S2). In contrast, the Foldseek TM score for each protein strongly influenced the likelihood of correct oligomeric state prediction, with proteins that had TM scores < 0.3 showing the lowest success rates (∼30-75% predicted correctly) regardless of filtering by pLDDT (Fig. 4B).

**Figure 4.**
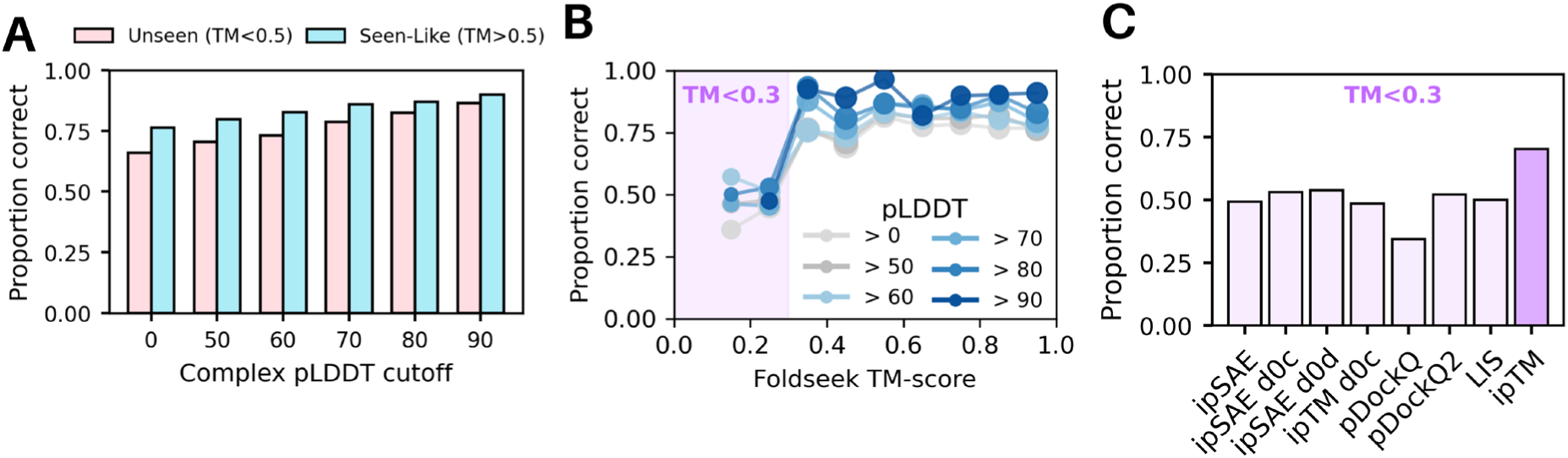
Oligomeric state prediction is less accurate for proteins that lack structural similarity to proteins in the AlphaFold training set. (A) Proportion of proteins predicted correctly by AF2-M from a set of approximately 500 “unseen” proteins (pink; no known structure, Foldseek TM score < 0.5 to any protein in the AlphaFold training set) and approximately 500 “seen-like” proteins (blue; no known structure, Foldseek TM score > 0.5 to at least one protein in the AlphaFold training set), filtered by different pLDDT cutoffs for the predicted oligomeric complexes. For monomeric proteins, the pLDDT value used corresponds to the folded oligomeric state with the highest mean ipTM. (B) Proportion of proteins predicted correctly as a function of Foldseek TM score. Data are filtered by the pLDDT of the predicted oligomeric complexes and shown as separate lines for each pLDDT cutoff. A purple square is added to highlight proteins with a TM score of < 0.3. Marker size is proportional to the number of proteins per bin. The number of proteins per bin is reported in Fig. S2E. (C) Proportion of proteins with Foldseek TM scores < 0.3 to any protein in the AlphaFold training set that were predicted correctly using different interface confidence metrics, including ipSAE, ipSAE d0 chn (ipSAE d0c), ipSAE d0 dom (ipSAE d0d), ipTM d0 chn (ipTM d0c), pDockQ, pDockQ2, LIS, and ipTM.

Recent studies have suggested that ipTM alone may be insufficient for assessing the quality of AlphaFold-predicted complexes and have proposed alternative interface-focused metrics such as ipSAE, ipSAE d0 chn, ipSAE d0 dom, ipTM d0 chn ^16^, pDockQ^17^, pDockQ2^18^, and LIS^19^. To determine whether these metrics improved oligomeric state prediction for structurally novel proteins, we evaluated prediction accuracy using each metric for proteins with TM scores < 0.3. However, none of the alternative metrics substantially outperformed ipTM for this subset (Fig. 4C). Similarly, ipTM also outperformed other metrics for identifying the correct oligomeric state for our full datasets of “Unseen” and “Seen-like” proteins (Fig. S3A-B). Together, these results indicate that proteins lacking structural similarity to the AlphaFold training set are intrinsically more difficult to classify correctly, and that this limitation is not overcome by current interface confidence metrics. Notably, ipTM also performed best for both unseen and seen-like proteins, even with filtering for pLDDT (Fig. S3).

While availability of the source code for AF2-M allowed us to generate structural predictions for many proteins without restriction, we wondered whether the online AlphaFold3 server which provides an accessible user interface to predict structural complexes could also be used to predict oligomeric states. To assess this, we carried out structural predictions of the 40 proteins as dimers, trimers, tetramers and pentamers as above using the default AF3 parameters of 10 recycles, 5 structures.

As with AF2-M, the ipTM scores of predicted oligomer structures can distinguish between proteins predicted in the annotated state versus an incorrect oligomeric state (Fig. 5A), as well as between monomeric and multimeric proteins (Fig. 5B). By optimising for the monomeric cutoff value to 0.24 as we did when using AF2-M, 33/40 of the proteins in our test set were correctly assigned (Fig. 4C, Fig. S4-5).

**Figure 5.**
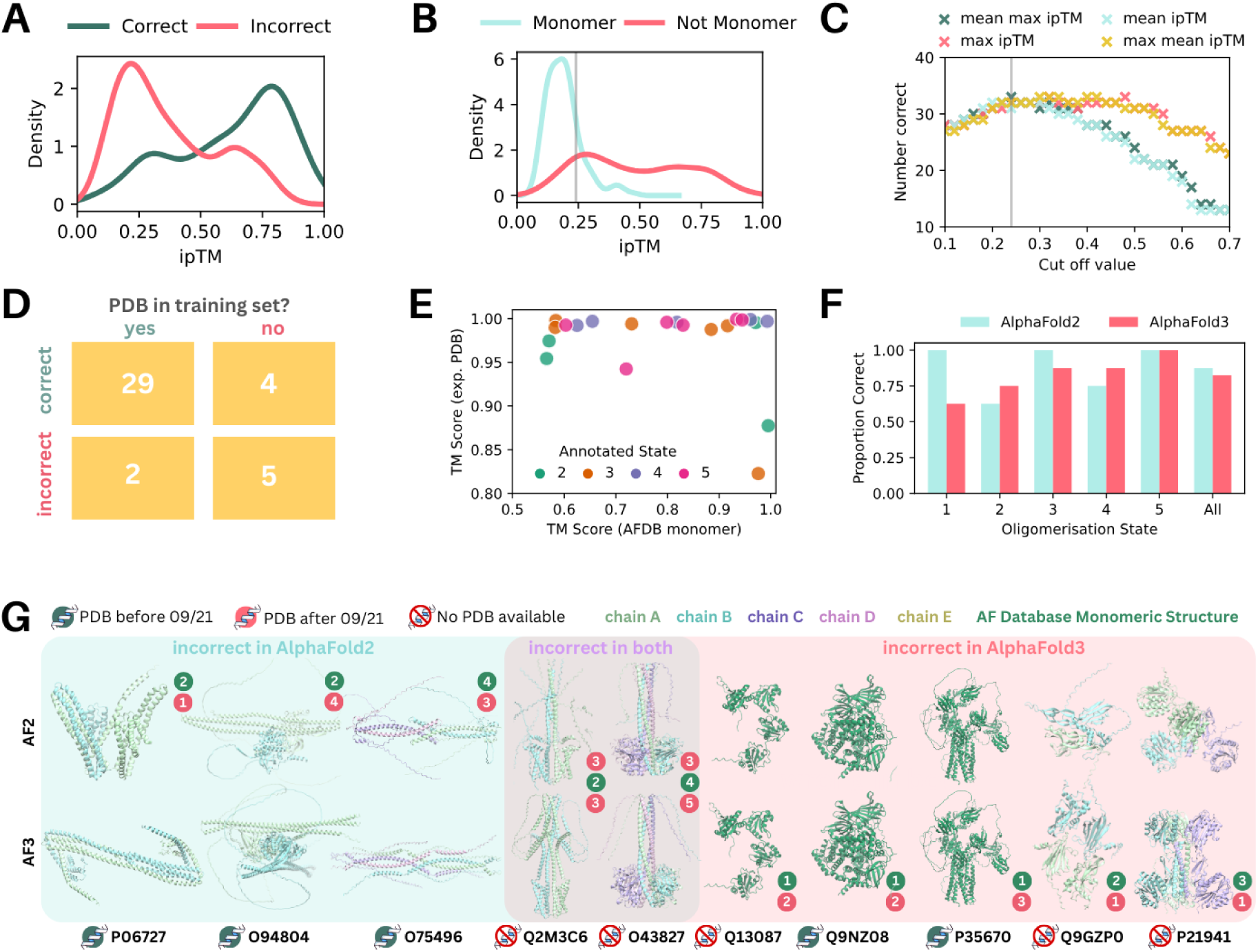
ipTM scores from the AlphaFold3 online server also accurately predict oligomeric states. (A) Probability distribution plot of ipTM scores for 32 known multimeric proteins predicted in the correct (pink) and incorrect (green) oligomeric states. (B) Probability distribution plot of ipTM scores from AF3 for 8 known monomeric proteins predicted with 2,3,4,5 subunits (blue) and 32 known multimeric proteins predicted with 2,3,4,5 subunits (pink). The grey line indicates the cut-off ipTM value of 0.24. (C) Number of correctly assigned oligomeric states (out of 40) vs cut-off value selected for 4 different methods to distinguish between monomeric and multi-subunit proteins. The grey line indicates the optimal cut-off ipTM value of 0.24. (D) Two-way table showing the distribution of proteins with oligomeric states predicted correctly and incorrectly based on whether an experimentally resolved structure exists within the training set. (E) TM score between the top ranked AlphaFold3 predicted structure and experimentally resolved structure (if available) plotted against the TM score between the top ranked AlphaFold3 predicted structure and the monomeric structure from the AFDB, coloured by annotated oligomerisation state. (F) Comparison of AlphaFold2 and AlphaFold3 prediction accuracy by annotated oligomerisation state. (G) Snapshots of the top ranked structural predictions generated by AF2-M (top row) and AF3 (bottom row) in the correct oligomeric state for proteins that were assigned to an incorrect oligomeric state by AF2-M, AF3, or both AF2-M and AF3 (ribbon, coloured by chain). For monomeric proteins, the AFDB monomeric structure is shown in dark green. The presence of an experimentally resolved structure in the PDB or AlphaFold2 training set is indicated with an icon near each UniProt ID. The correct oligomerisation state is indicated by each structure in green and the incorrect prediction by the associated method in red.

As AF3 uses PDB templates by default where available, we find most proteins (29/31) where a structure was present in the training set is predicted in the correct oligomeric state (Fig. 5D). In contrast, 5/9 structures which are not present in the training set were predicted in the incorrect oligomeric state.

Similar to AF2-M, the majority of predicted oligomeric structures had high TM scores to the experimentally resolved structures where available, but a lower correlation of TM scores to the predicted monomeric structures (Fig. 5E). Similar TM scores were obtained for the same proteins across AF2-M and AF3, and proteins which were predicted with lower TM scores using AF2-M were also less like the experimental structures when predicted using AF3.

Compared to AlphaFold2, AlphaFold3 assigned more dimeric and tetrameric proteins correctly (Fig. 5F), although the significance of this is hard to assess given the size of the data set. Structures predicted in an incorrect oligomeric state by AF2-M but had a PDB available were predicted in the correct oligomeric state by AF3 (Fig. 5G). Interestingly, 5/7 of the incorrect state assignments using AF3 was due to difficulty distinguishing between monomeric and homooligomeric proteins. Three monomeric proteins were predicted to form higher order oligomers (2 dimers, 1 trimer) and one dimeric and one trimeric protein was incorrectly assigned as monomeric.

Next, we sought to apply this technique of oligomerisation state and structure prediction to membrane proteins. As many pore forming membrane proteins are multimeric, their oligomeric state is crucial for understanding their physiology. We curated a set of 14 membrane proteins with a diversity of known oligomeric states (monomeric to heptameric) of interest in our lab and used these to assess AF2-M’s ability to correctly predict their oligomeric states (Fig. 6A, Table S3).

**Figure 6.**
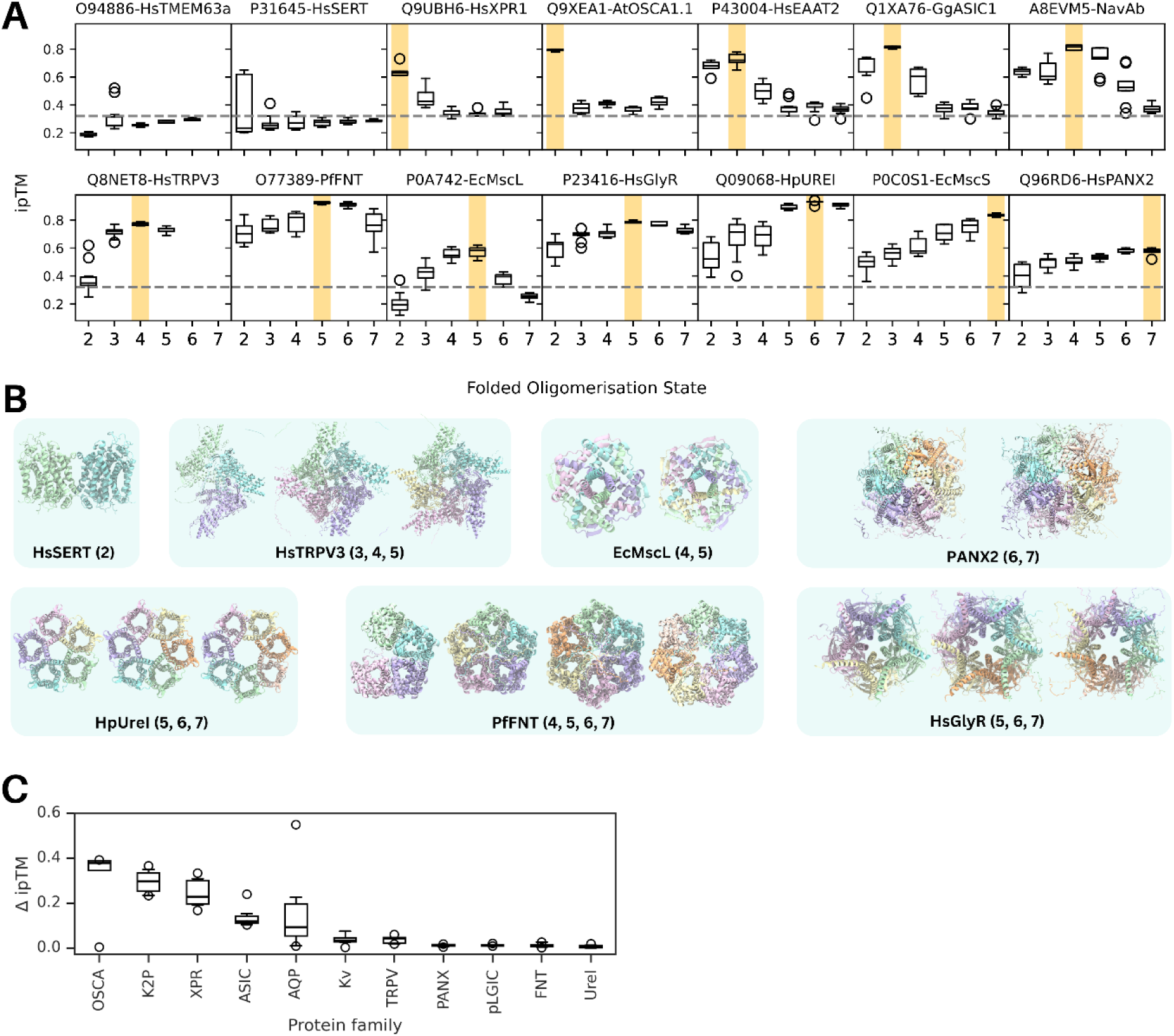
Application to membrane proteins. (A) Box plots showing individual ipTM scores for different folded oligomeric states (2-7) across a curated set of 14 membrane proteins with the correct oligomeric state highlighted in yellow. The dashed line indicates the previously determined AF2-M cut-off ipTM value of 0.32. UniProt IDs and common protein names are shown. (B) Structural predictions of select membrane proteins in various oligomeric states. Protein names and oligomeric states shown are given below each set of structures. (C) Box plots (left) and density plots (right) showing the difference in ipTM (ΔipTM) between the top two folded oligomeric states with the highest mean ipTM for each protein, grouped by family. Each dot in the box plot represents a single protein from the OSCA, K2P, XPR, ASIC, AQP, Kv, TRPV, PANX, pLGIC, FNT, or UreI families.

In this curated set, all proteins which form higher order oligomers (dimers and above) were predicted in the correct oligomeric states. However, two proteins which we expected to be assigned as monomeric (HsTMEM63a, UniProt ID: O94886 and HsSERT, UniProt ID: P31645) were assigned as trimers and dimers respectively using our monomeric cut-off value of 0.32. Interestingly, we note that for HsSERT, the top ranked structures of the dimeric form of the protein (as shown in Fig 6B) have the highest ipTM score (Fig 6A). Dimerisation and formation of higher order oligomers for HsSERT and other SLC6 members have been previously reported, hinting that the oligomerisation state predicted here could be biological relevant.^20^ Additionally, the top ranked predicted dimeric structure of HsSERT resembles one dimer conformation sampled in previously conducted molecular dynamics simulations probing HsSERT self-assembly.^21^

The transient receptor potential vanilloid-3 channel TRPV3 (UniProt ID: Q8NET8) was correctly predicted to be tetrameric. In addition, we noticed the generally high ipTM scores and low ipTM range of structures predicted in the trimeric and pentameric states, which both had ipTM scores only slightly lower than the tetramers. The predicted trimeric structures resembled the tetramers with one missing subunit (Fig 5B), highlighting a case where visual examination is necessary to understand whether a predicted oligomeric state is plausible. Interestingly, however, the pentameric structures resembled the cryo-EM structure of TRPV3 recently elucidated (after the AF2.3 training date cutoff) as a pentamer, thought to be a transient state of the protein with a dilated pore.^22^

Similarly, we observed that the mechanosensitive channel of large conductance MscL, whose structure has been determined in both the tetrameric and pentameric states^23,24^, had similar ipTM score distributions for these two states. However, we note in this case that both structures were determined prior to the cut-off date for training AF2. Finally, structural predictions of Pannexin 2 suggest that the hexameric and heptameric oligomerisation states are both probable, as they have similar high ipTM scores. Our results are consistent with a recent report that the closely related pannexin 1 channels are active in both the hexameric and heptameric states.^25^ This result was surprising given that all structures of pannexins thus far have been resolved as heptamers.^26–30^

Interestingly, the ipTM score distributions and structural predictions for HpUreI (a proton-gated urea channel in *H. pylori*)^31^, PfFNT (a lactate/proton transporter in *P. falciparum*)^32,33^ and HsGlyR (the human glycine receptor)^34^ suggest that these proteins may also oligomerise with a different number of subunits in addition to their currently known annotated states.

As HsTRPV3, HsPANX2, PfFNT, HsGlyR and HPUreI all had at least two oligomeric states with similar ipTM values, we extended our investigations with AF2-M to additional proteins from the same families (TRPV, PANX, FNT, pLGIC, and UreI, respectively) to see if a similar trend would be observed in the range of ipTM values (Fig. 6C, Table S4). For comparison, we also folded various proteins from the OSCA, XPR, ASIC, AQP, K2P, and Kv families. For each folded protein, the difference in ipTM (ΔipTM) between the top two folded oligomeric states with the highest mean ipTM was calculated. Interestingly, proteins from the TRPV, PANX, FNT, pLGIC, UreI, and Kv families generally had lower ΔipTM values than those from the OSCA, XPR, ASIC, AQP, and K2P families. While HsTRPV3 is known to adopt different oligomeric states and HsPANX2 has been hypothesised to adopt different oligomeric states, and both proteins have low ΔipTM values, it is plausible that comparing ΔipTM between folded oligomeric states may suggest if a protein is likely to adopt different oligomeric states. However, we note the presence of apparent false positives, including the case of the Kv family, which show low ΔipTM values between tetramers and pentamers. Although one study has reported a pentameric structure of the soluble T1 domain of Kv2.1^35^, full length Kv channels have only been resolved as tetramers.

## Discussion

Many proteins undergo oligomerisation to form their biologically relevant states, and predicting both stoichiometry and structure is critical for understanding protein function and regulation. Here, we show that AlphaFold2-Multimer and AlphaFold3 can be used to predict oligomeric states for a subset of proteins with high confidence, while also revealing clear limitations when predictions are extended to larger and more structurally diverse datasets. Using both a curated benchmark set and a large-scale collection of over 1,000 proteins, we identify the conditions under which oligomeric state prediction is reliable, and those under which it remains challenging. As the benchmark dataset contained only human proteins, whereas the large-scale expanded dataset included proteins from multiple species, indicating that AlphaFold-based oligomeric state prediction may be largely species agnostic.

We optimise the computational efficiency of our prediction method, showing that 3 structures predicted with 10 recycles using AF2-M and the default prediction parameters available on the AF3 online server are sufficient to derive correct oligomeric states for 35/40 and 33/40 of the proteins in our test set respectively. The small number of proteins in our test set and differences in parameters used means that it is not possible to say here whether AF2-M or AF3 is more likely to predict the correct oligomeric state. However, it appears that within this set of proteins, proteins with generally high pLDDT scores and without abnormally long helices in the AFDB structure have their oligomeric states correctly predicted by AF2-M. It is less clear what determines whether a protein will be predicted in the correct oligomeric state by AF3, although it seems to perform less well for proteins without an available structure in the PDB and has some trouble distinguishing between monomeric and non-monomeric proteins. We note that in 30/31 cases, if both AF2-M and AF3 predict the same oligomeric state for a given protein, then this is the annotated state, suggesting that use of AF2-M and AF3 concurrently could increase confidence in the oligomeric state prediction.

While average monomeric pLDDT is a strong indicator of oligomeric state prediction accuracy in the small, curated dataset, analysis of the larger dataset of 1,006 proteins demonstrates that high pLDDT alone is not sufficient to guarantee correct oligomeric state prediction. This limitation is particularly apparent for proteins that lack similarity to any structures present in the AlphaFold training set, highlighting a challenge for extending oligomeric state prediction to proteins that are more structurally diverse and evolutionarily distant.

To determine whether this limitation arises from the confidence metrics used rather than the underlying structural predictions, we examined alternative interface-based scoring approaches. The ipTM metric used to identify the most likely oligomeric state is reported by default for each structure in both AF2-M and AF3 and performs better than pLDDT values which are not expected to be strongly influenced by the interchain contacts. However, it has been noted that the ipTM scores can be impacted by the presence of disordered protein regions.^16,36,37^ The effect of these regions can be reduced by removing them if they are known not to be involved in the interaction, restricting calculations to only residues near the interaction site,^38^ or to interchain residue pairs that have well predicted aligned error distances.^16^ While these approaches may improve ipTM-based assessments in specific cases, our large-scale benchmarking indicates that more advanced interface confidence metrics, including ipSAE^16^, pDockQ^17^, pDockQ2^18^, and LIS^19^, do not substantially improve oligomeric state prediction for proteins for which AlphaFold has no close structural reference. This suggests that the primary limitation lies not in the choice of interface metric, but in the absence of sufficient evolutionary or structural context for AlphaFold to predict accurate inter-chain interactions.

Despite these limitations, ipTM distributions can still provide valuable information about protein oligomeric states. For several membrane proteins which can be physiologically found in different oligomeric states (TRPV3, MscL and Pannexins), this information can be captured in the ipTM distributions. This suggests the value of this approach for identifying proteins where multiple physiologically relevant oligomerization states may be plausible and warrant further investigation.

We further extended this analysis to proteins from the OSCA, K2P, XPR, ASIC, AQP, Kv, TRPV, PANX, pLGIC, FNT, and UreI families. Proteins belonging to families containing at least one member previously reported to adopt multiple oligomeric states exhibited small differences in ipTM values between two or more folded oligomeric states. This pattern suggests that low ΔipTM between folded oligomeric states may be able to suggest proteins capable of adopting multiple biologically relevant oligomeric forms. However, we note that this method is not completely accurate, as members of the Kv family also showed low ΔipTM values despite full length channels not having been reported to adopt different oligomeric states.

These findings also highlight a limitation of relying on existing database annotations for knowledge of protein oligomeric state. For two membrane proteins we examined (HsXPR1^39–41^ and NavAb^42^), oligomeric states determined via structural biology were not annotated in the UniProt database. Additionally, for PANX1, which has recently been shown to be functional as both hexamers and heptamers, only the heptameric state is annotated, suggesting that interaction annotations in UniProt may sometimes provide an incomplete picture of physiologically relevant oligomerisation.

Despite its benefits, it is also key to consider the computational feasibility of this approach. The high computational cost of modelling four or more copies of larger proteins may prohibit this technique from being easily usable for proteins where one subunit contains more than 1000 amino acids. Additionally, due to computational constraints, we did not consider hexameric and higher-order assemblies in either the curated benchmark set of 40 proteins or the expanded dataset of 1,006 proteins, with the exception of a small number of hexameric and heptameric membrane proteins shown in Fig. 5.

Future studies will therefore be needed to assess the performance of this technique for higher-order oligomeric complexes to determine whether this approach generalises beyond pentamers, as it may be harder to distinguish higher order pentamers with more similar angles between subunits. Further extensions of this work may also include investigating whether this technique could provide insight into stoichiometric ratios for heteromeric complexes in instances where the identity of interacting proteins is known but the number of each subunit is not.

Beyond oligomeric state prediction, an additional advantage of using AlphaFold is that it provides structural models of proteins in their predicted assemblies, which may enable further functional insight. To increase confidence in these models, plausible oligomeric structures could be combined with short, physics-based molecular dynamics simulations to assess their stability in solution or membrane environments, alongside experimental validation.

Ultimately, AlphaFold2-Multimer and AlphaFold3 represent powerful tools for predicting oligomeric states and generating structural models, particularly for proteins with fold-level similarity to known structures. However, our results demonstrate that accurate oligomeric state prediction remains challenging for structurally novel proteins, even when predicted structures are of high confidence and evaluated using advanced interface metrics. Together, these findings define both the promise and the current limitations of AlphaFold-based oligomeric state prediction and highlight the continued importance of experimental validation for proteins for which there are currently no structurally similar representatives in the AlphaFold training set.

## Methods

### Selection of oligomeric protein sequences

The DeepSub database (https://github.com/tibbdc/DeepSub/blob/main/DATA/Dataset_0724_new.csv) containing proteins from the UniProt database annotated with experimentally determined oligomeric states was used to randomly select 40 human proteins (5 each of known monomeric, homodimeric, homotrimeric, homotetrameric and homopentameric proteins), detailed in Table S1.^6^

### Structural prediction using AlphaFold2 Multimer

Selected sequences were extracted from the UniProt Database and used to generate multiple sequence alignments for structure prediction by ColabFold implementing AlphaFold-Multimer v.2.3.^43,44^ 20 structures containing 2,3,4,5 copies of each protein were predicted with 3 recycles without subsequent relaxation nor modification to the multiple sequence alignment. To optimize computational efficiency, structural predictions were also carried out using all permutations of 1,5,10,20 structures and 0,1,2,3 recycles. For the 5 proteins with incorrect oligomeric state predictions, we carried out additional structural predictions using 5 and 10 recycles (generating 20 structures in each case). For two large proteins within our dataset (UniProt IDs: P35670, P533396), we were unable to predict their structures as tetramers/pentamers due to computational restrictions.

To extend our assessment of the accuracy of using AF2-M to predict the oligomeric states of proteins, we randomly selected 1,006 proteins from the DeepSub database for which the oligomeric state was known but no structures were available. To ensure that none of the test proteins had homologs represented in the AlphaFold2 (AF2) training set, proteins from the DeepSub database were first screened to confirm the absence of experimentally determined structures. Monomeric AFDB models for the remaining proteins were downloaded, and structural similarity searches were performed against all PDB entries using FoldSeek. A TM-score threshold of 0.5 was used to identify and exclude proteins with detectable structural similarity.

These proteins were divided almost equally into two categories “seen-like” and “unseen”, based on the Foldseek determined TM score between the AFDB monomeric structure and experimentally determined structures in the AlphaFold training set. “Seen-like” proteins had at least one homologous structure with a Foldseek TM score > 0.5, whereas “unseen” proteins showed no detectable similarity to any PDB structure (TM score < 0.5), corresponding to proteins for which AlphaFold has no close structural reference.

Both the “seen-like” and “unseen” groups contained at least 40 monomers and approximately 100 dimers, trimers, tetramers, and pentamers. Each protein was then modelled as a dimer, trimer, tetramer, and pentamer using AF2 with 3 recycles, generating 20 structural models for each prediction.

To evaluate oligomeric-state predictions, we compared raw ipTM scores from AF2 with a set of advanced interface-quality metrics. This included the ipSAE, ipSAE-d0chn, ipSAE-d0dom, ipTM-d0chn^16^, pDockQ^17^, pDockQ2^18^, and LIS scores^19^. These metrics were computed using code obtained from the Dunbrack lab’s repository (https://github.com/dunbracklab/IPSAE).

AF2-M was used to fold an additional 98 proteins from the OSCA, K2P, XPR, ASIC, AQP, Kv, TRPV, PANX, pLGIC, FNT, and UreI families as dimers, trimers, tetramers, pentamers, hexamers, and heptamers. All predictions were run using 3 recycles and 20 structures. For each protein, the top two folded oligomeric states with the highest mean ipTM were determined. A mean ipTM value was calculated for the second highest state and was subtracted from each ipTM value from the highest state, and these values were then averaged to calculate the mean ΔipTM per protein.

### Structural prediction using AlphaFold3

The AlphaFold3 online server was used to predict multimeric complexes (2,3,4,5 copies) of each protein, using sequences extracted from the UniProt Database.^11^ By default, 10 recycles were performed and 5 structures generated. As above, for two large proteins (UniProt IDs: P35670, P533396), we were unable to predict their structures as tetramers/pentamers due to computational restrictions.

### Analysis of structures

Structures and output files produced by AlphaFold2-Multimer and AlphaFold3 were analysed using python scripts written using the pandas, scipy, matplotlib, MDAnalysis and MDTraj libraries.^45–49^ Structures were visualised and figures produced using Visual Molecular Dynamics (VMD).^50^

For each protein, the predicted monomeric structure from the AFDB and the top ranked structure produced for the correct oligomeric state were oriented along the Z principal axis using the GMX editconf princ function within GROMACS (v.2023).^51^ For oligomeric structures predicted using AlphaFold2-Multimer, the top ranked structure was chosen from those generated from 3 recycles and 20 structures. The α-carbon atoms of the monomeric structure and the first chain of the predicted oligomeric structure were aligned within VMD to generate the structural images.

The average monomeric pLDDT for each protein was calculated using each associated structure in the AFDB by averaging the per-residue pLDDT for all residues in each structure.

The secondary structure was calculated using the DSSP algorithm implemented in MDTraj.^49,52^ For each structure, the length of the longest helix was assigned to be the longest continuous stretch of residues calculated to have a helical (‘H’) secondary structure.

The template modelling (TM) score for the structures of each protein predicted in the correct oligomeric state were calculated following sequence and structural alignment to the predicted monomeric structure in the AFDB and, where available, to valid experimentally resolved structures using US-align.^53^ All experimentally resolved structures associated with the UniProt ID of each structure were assessed for validity. Monomeric structures were removed from comparison to our oligomeric predictions. Structures with < 95% sequence similarity between the UniProt sequence and PDB sequence were discarded, as were structures containing only fragments (< 50% of the UniProt sequence) of the protein. Using options –mm 1 and –ter 0, US-Align was used to calculate the TM score after alignment of the two multi-chain oligomeric structures. As some proteins of interest were resolved with accessory proteins and additional chains were present in some PDBs, structures were manually inspected to identify the correct chains for comparison where necessary. For the 5 proteins with a large number (> 10) of eligible PDBs (Table S3), if a PDB with the correct number of chains yielded a TM score above 0.95, we did not calculate further TM scores for PDBs resolved with additional chains. TM scores to the monomeric structure were also calculated using US-Align, normalizing by the length of the monomeric protein.

## Contributions

Conceptualisation: Y.L., C.W., B.C.

AlphaFold2 Predictions: C.W., Y.L.

AlphaFold3 Predictions: Y.L., C.W.

Benchmarking: C.W., Y.L.

Dataset curation: C.W., Y.L.

Analysis: Y.L., C.W.

Figures: Y.L., C.W.

Manuscript writing: Y.L., C.W., B.C.

Funding: B.C., Y.L.

## Acknowledgements

This research was supported by services and resources provided by the National Computational Infrastructure (NCI), which was funded by the Australian Government. We acknowledge funding from the Australian Research Council (DP200100860), and National Health and Medical Research Council (APP2020565). C.W. acknowledges support from an Australian Government Research Training Program (RTP) Ph.D. Scholarship.

## Supporting Information

**Table S1.**
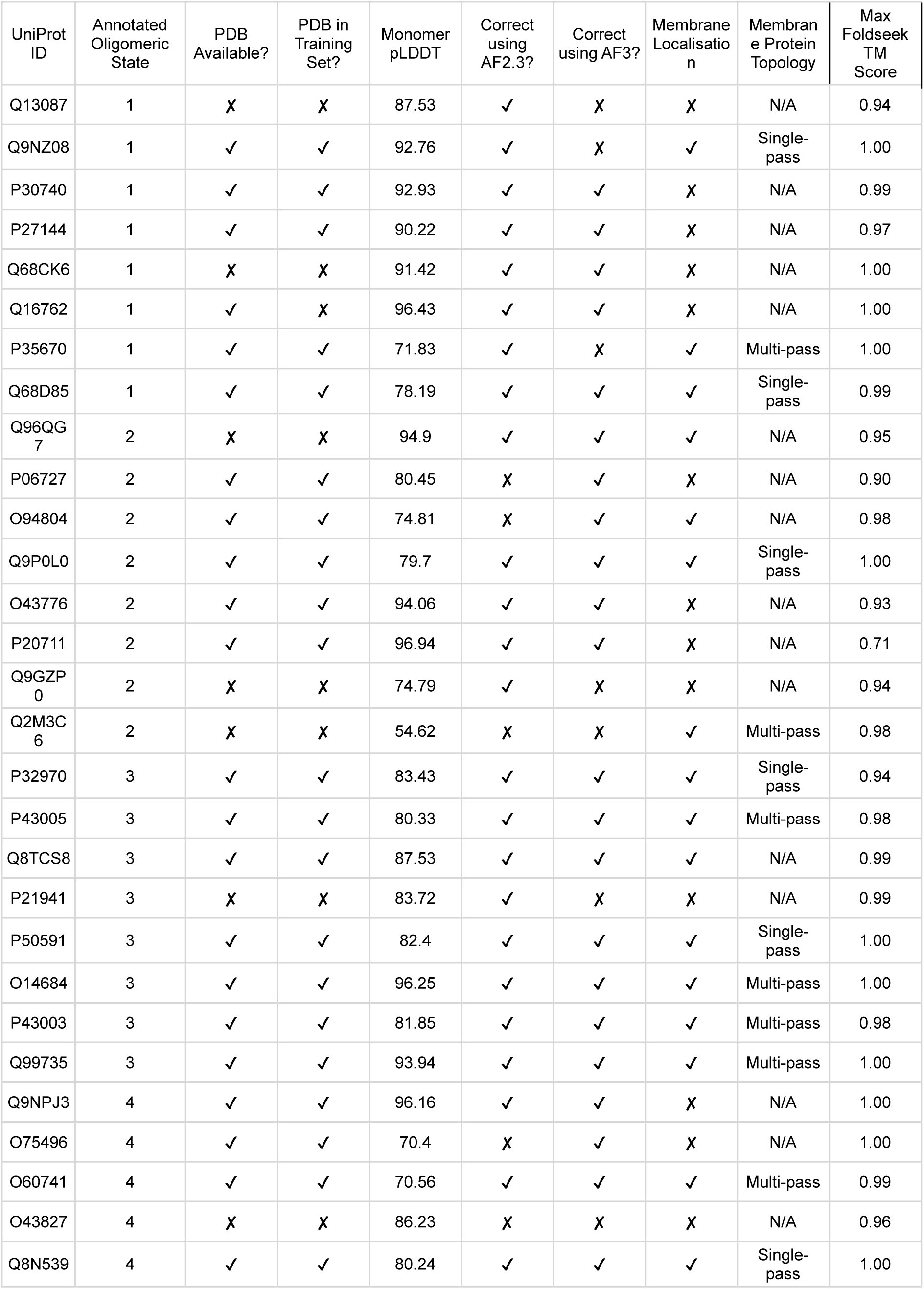
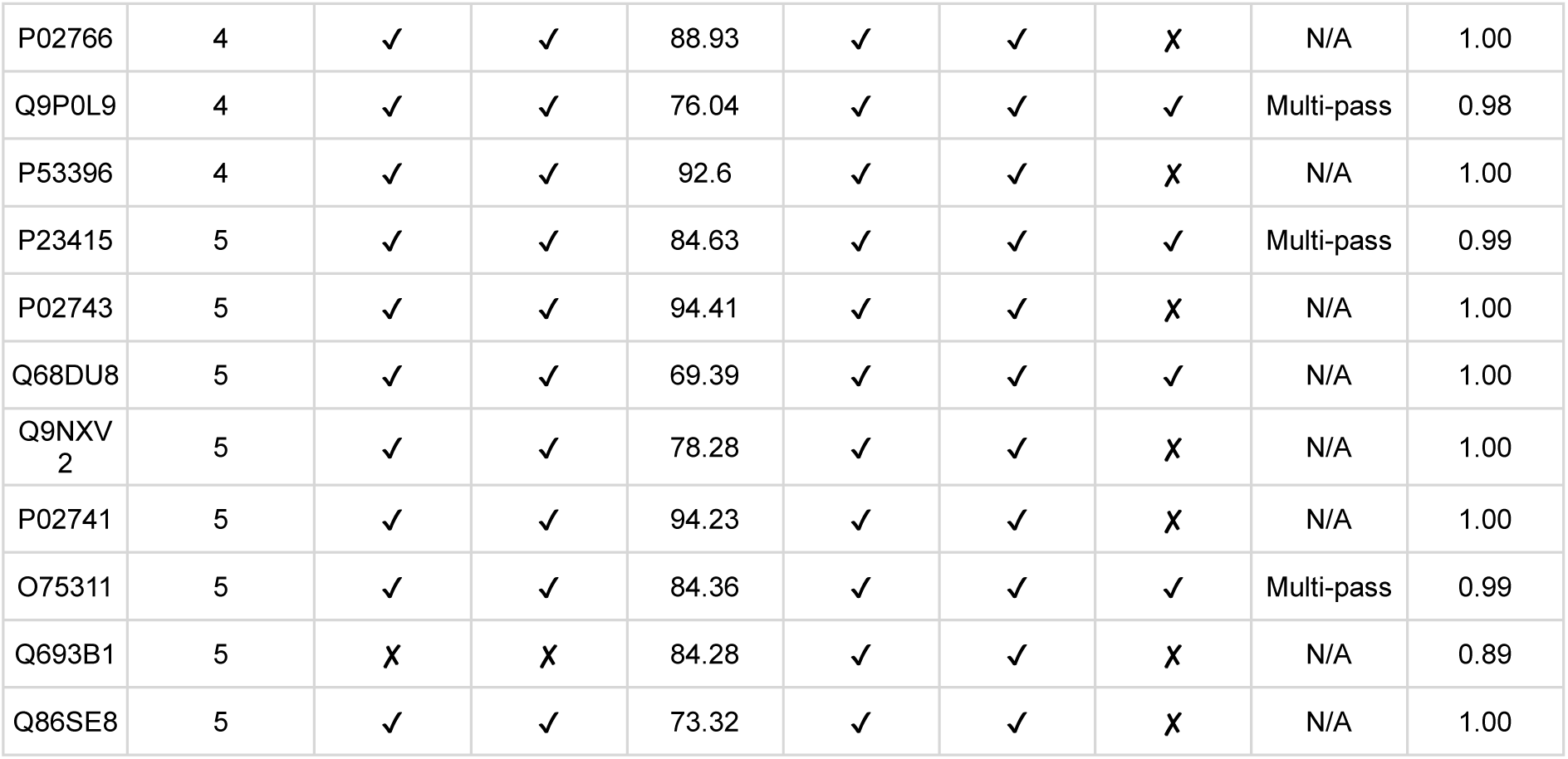
– Protein Set.

| UniProt ID | Annotated Oligomeric State | PDB Available? | PDB in Training Set? | Monomer pLDDT | Correct using AF2.3? | Correct using AF3? | Membrane Localisation | Membrane Protein Topology | Max Foldseek TM Score |
| --- | --- | --- | --- | --- | --- | --- | --- | --- | --- |
| Q13087 | 1 | ✗ | ✗ | 87.53 | ✓ | ✗ | ✗ | N/A | 0.94 |
| Q9NZ08 | 1 | ✓ | ✓ | 92.76 | ✓ | ✗ | ✓ | Single-pass | 1.00 |
| P30740 | 1 | ✓ | ✓ | 92.93 | ✓ | ✓ | ✗ | N/A | 0.99 |
| P27144 | 1 | ✓ | ✓ | 90.22 | ✓ | ✓ | ✗ | N/A | 0.97 |
| Q68CK6 | 1 | ✗ | ✗ | 91.42 | ✓ | ✓ | ✗ | N/A | 1.00 |
| Q16762 | 1 | ✓ | ✗ | 96.43 | ✓ | ✓ | ✗ | N/A | 1.00 |
| P35670 | 1 | ✓ | ✓ | 71.83 | ✓ | ✗ | ✓ | Multi-pass | 1.00 |
| Q68D85 | 1 | ✓ | ✓ | 78.19 | ✓ | ✓ | ✓ | Single-pass | 0.99 |
| Q96QG7 | 2 | ✗ | ✗ | 94.9 | ✓ | ✓ | ✓ | N/A | 0.95 |
| P06727 | 2 | ✓ | ✓ | 80.45 | ✗ | ✓ | ✗ | N/A | 0.90 |
| O94804 | 2 | ✓ | ✓ | 74.81 | ✗ | ✓ | ✓ | N/A | 0.98 |
| Q9P0L0 | 2 | ✓ | ✓ | 79.7 | ✓ | ✓ | ✓ | Single-pass | 1.00 |
| O43776 | 2 | ✓ | ✓ | 94.06 | ✓ | ✓ | ✗ | N/A | 0.93 |
| P20711 | 2 | ✓ | ✓ | 96.94 | ✓ | ✓ | ✗ | N/A | 0.71 |
| Q9GZP0 | 2 | ✗ | ✗ | 74.79 | ✓ | ✗ | ✗ | N/A | 0.94 |
| Q2M3C6 | 2 | ✗ | ✗ | 54.62 | ✗ | ✗ | ✓ | Multi-pass | 0.98 |
| P32970 | 3 | ✓ | ✓ | 83.43 | ✓ | ✓ | ✓ | Single-pass | 0.94 |
| P43005 | 3 | ✓ | ✓ | 80.33 | ✓ | ✓ | ✓ | Multi-pass | 0.98 |
| Q8TCS8 | 3 | ✓ | ✓ | 87.53 | ✓ | ✓ | ✓ | N/A | 0.99 |
| P21941 | 3 | ✗ | ✗ | 83.72 | ✓ | ✗ | ✗ | N/A | 0.99 |
| P50591 | 3 | ✓ | ✓ | 82.4 | ✓ | ✓ | ✓ | Single-pass | 1.00 |
| O14684 | 3 | ✓ | ✓ | 96.25 | ✓ | ✓ | ✓ | Multi-pass | 1.00 |
| P43003 | 3 | ✓ | ✓ | 81.85 | ✓ | ✓ | ✓ | Multi-pass | 0.98 |
| Q99735 | 3 | ✓ | ✓ | 93.94 | ✓ | ✓ | ✓ | Multi-pass | 1.00 |
| Q9NPJ3 | 4 | ✓ | ✓ | 96.16 | ✓ | ✓ | ✗ | N/A | 1.00 |
| O75496 | 4 | ✓ | ✓ | 70.4 | ✗ | ✓ | ✗ | N/A | 1.00 |
| O60741 | 4 | ✓ | ✓ | 70.56 | ✓ | ✓ | ✓ | Multi-pass | 0.99 |
| O43827 | 4 | ✗ | ✗ | 86.23 | ✗ | ✗ | ✗ | N/A | 0.96 |
| Q8N539 | 4 | ✓ | ✓ | 80.24 | ✓ | ✓ | ✓ | Single-pass | 1.00 |
| P02766 | 4 | ✓ | ✓ | 88.93 | ✓ | ✓ | ✗ | N/A | 1.00 |
| Q9P0L9 | 4 | ✓ | ✓ | 76.04 | ✓ | ✓ | ✓ | Multi-pass | 0.98 |
| P53396 | 4 | ✓ | ✓ | 92.6 | ✓ | ✓ | ✗ | N/A | 1.00 |
| P23415 | 5 | ✓ | ✓ | 84.63 | ✓ | ✓ | ✓ | Multi-pass | 0.99 |
| P02743 | 5 | ✓ | ✓ | 94.41 | ✓ | ✓ | ✗ | N/A | 1.00 |
| Q68DU8 | 5 | ✓ | ✓ | 69.39 | ✓ | ✓ | ✓ | N/A | 1.00 |
| Q9NXV<br>2 | 5 | ✓ | ✓ | 78.28 | ✓ | ✓ | ✗ | N/A | 1.00 |
| P02741 | 5 | ✓ | ✓ | 94.23 | ✓ | ✓ | ✗ | N/A | 1.00 |
| O75311 | 5 | ✓ | ✓ | 84.36 | ✓ | ✓ | ✓ | Multi-pass | 0.99 |
| Q693B1 | 5 | ✗ | ✗ | 84.28 | ✓ | ✓ | ✗ | N/A | 0.89 |
| Q86SE8 | 5 | ✓ | ✓ | 73.32 | ✓ | ✓ | ✗ | N/A | 1.00 |

**Table S2.**
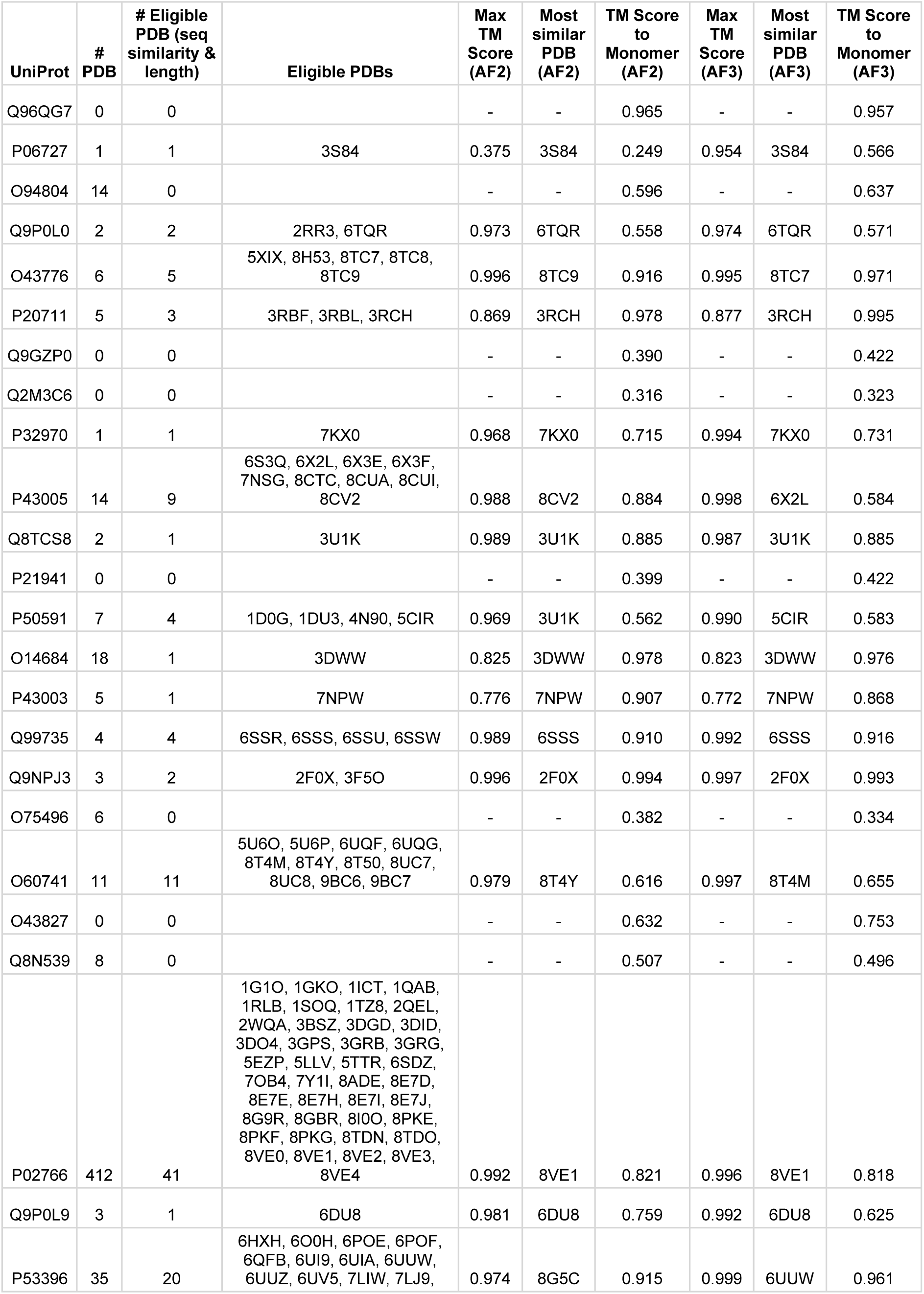
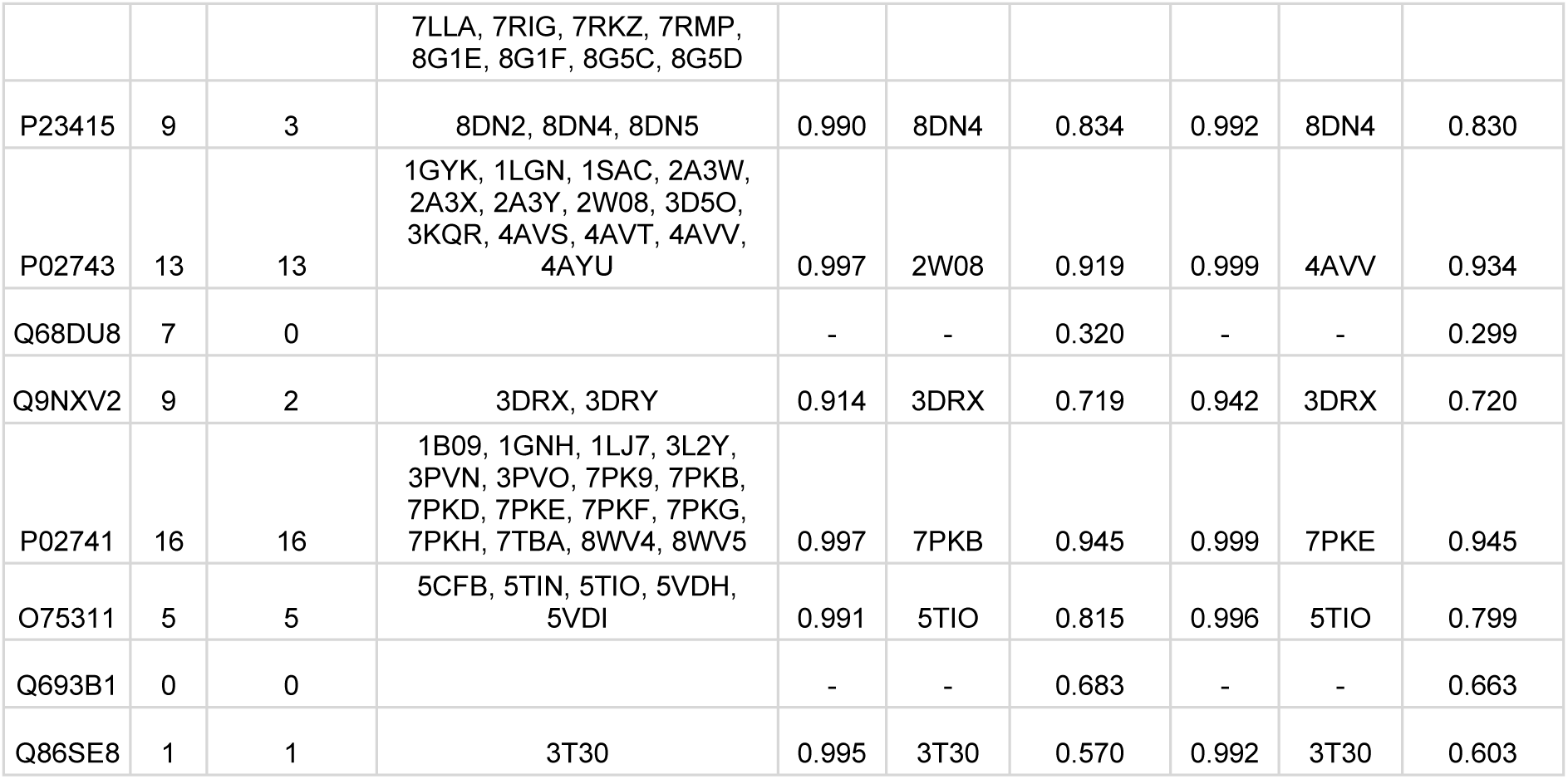
– Comparison of oligomer predictions to experimentally resolved structures and AF2 monomer.

**Table S3.** – Membrane Protein Set.

| UniProt ID | Annotated Oligomeric State | PDB Available? | PDB in Training Set? | Monomer pLDDT | Correct using AF2.3? | Membrane Localisation? | Membrane Protein Topology | Max Foldseek TM Score |
| --- | --- | --- | --- | --- | --- | --- | --- | --- |
| O94886 | 1 | ✓ | ✗ | 75.3 | ✗ | ✓ | Multi-pass | 0.86 |
| P31645 | 1 | ✓ | ✓ | 85.43 | ✗ | ✓ | Multi-pass | 0.99 |
| Q9UBH6 | 2 | ✓ | ✓ | 83.73 | ✓ | ✓ | Multi-pass | 0.90 |
| Q9XEA1 | 2 | ✓ | ✓ | 84.81 | ✓ | ✓ | Multi-pass | 0.99 |
| P43004 | 3 | ✓ | ✗ | 77.74 | ✓ | ✓ | Multi-pass | 0.97 |
| Q1XA76 | 3 | ✓ | ✓ | 83.42 | ✓ | ✓ | Multi-pass | 0.98 |
| A8EVM5 | 4 | ✓ | ✓ | 93.73 | ✓ | ✓ | Multi-pass | 1.00 |
| Q8NET8 | 4 | ✓ | ✓ | 76.93 | ✓ | ✓ | Multi-pass | 0.98 |
| O77389 | 5 | ✓ | ✓ | 91.42 | ✓ | ✓ | Multi-pass | 0.95 |
| P0A742 | 5 | ✓ | ✓ | 85.22 | ✓ | ✓ | Multi-pass | 0.57 |
| P23416 | 5 | ✓ | ✓ | 84.44 | ✓ | ✓ | Multi-pass | 0.99 |
| Q09068 | 6 | ✗ | ✗ | 90.27 | ✓ | ✓ | Multi-pass | 0.96 |
| P0C0S1 | 7 | ✓ | ✓ | 92.62 | ✓ | ✓ | Multi-pass | 0.95 |
| Q96RD6 | 7 | ✓ | ✗ | 58.15 | ✓ | ✓ | Multi-pass | 0.90 |

**Table S4.** – Extended Membrane Protein Set. See supplementary File

**Table S5.** – Unseen and Seen-like Protein Set. See supplementary file

**Figure S1.**
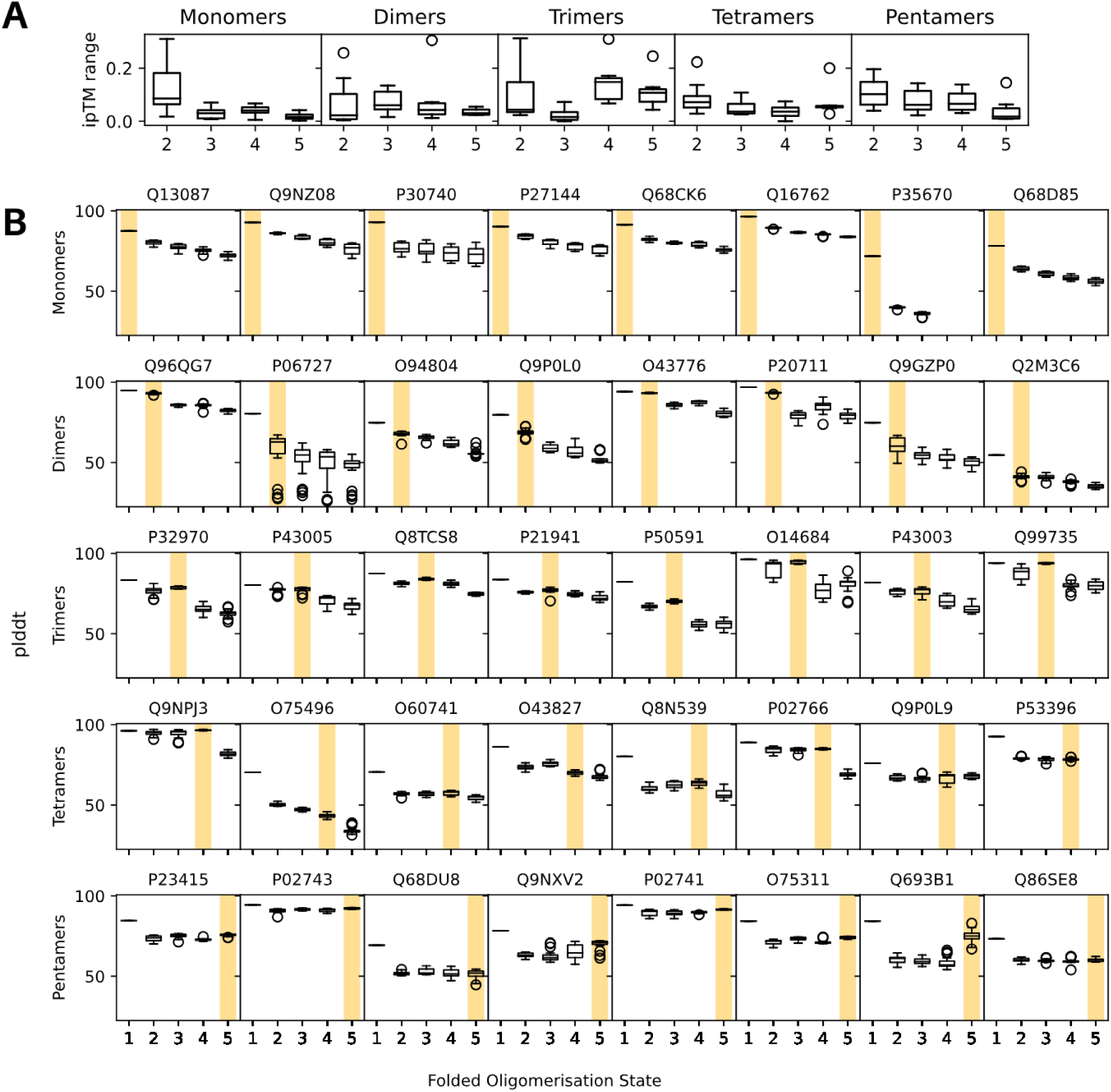
– ipTM range and pLDDT plots for AF2-M predictions. (a) ipTM range distributions for monomeric, dimeric, trimeric, tetrameric and pentameric proteins in different folded oligomeric states (b) Box plots showing pLDDT score distributions for each protein in different folded oligomeric states across the 40 test set proteins. The annotated oligomeric state highlighted in yellow.

**Figure S2.**
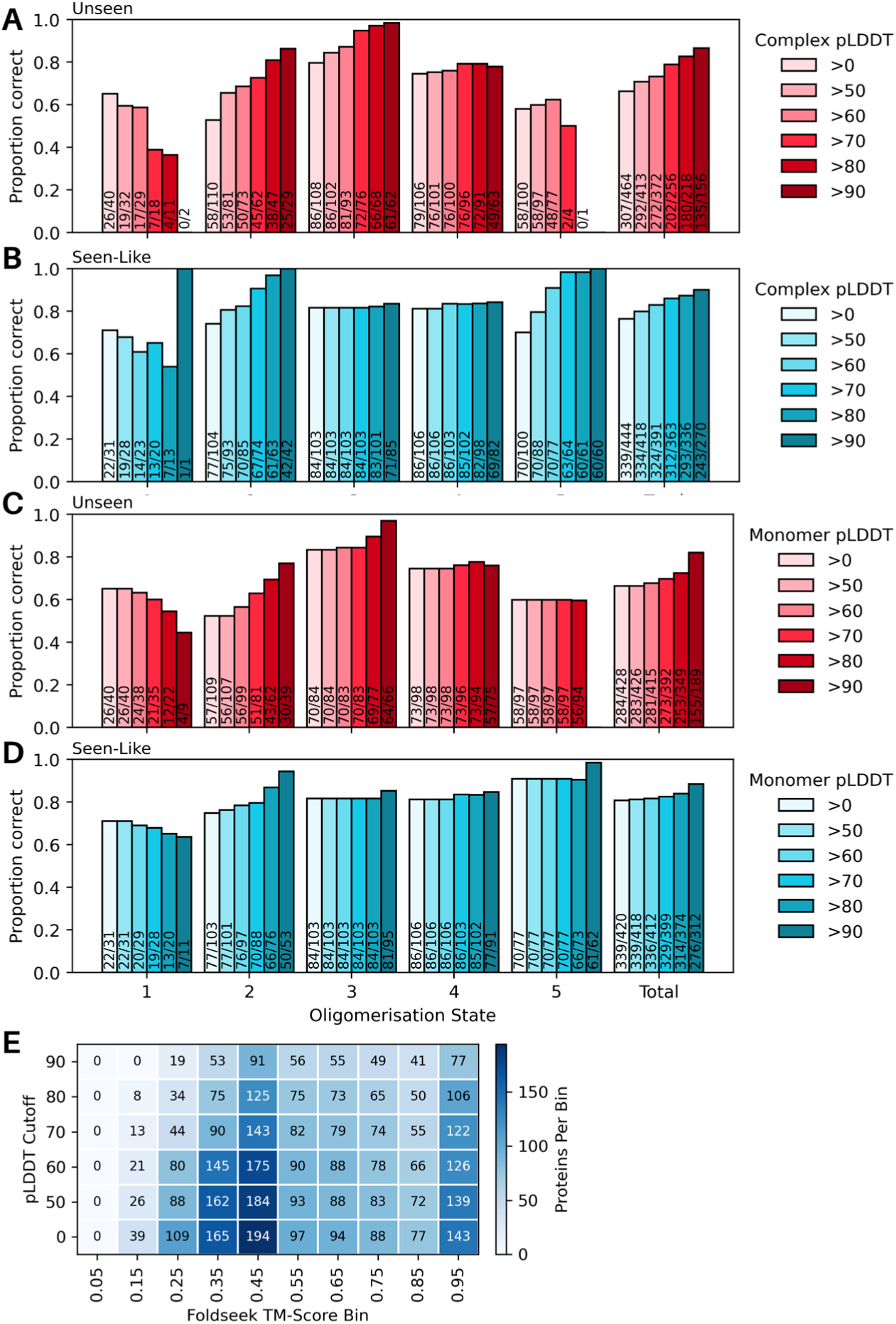
– Proportion of unseen and seen-like proteins predicted in the correct oligomeric state by AF2-M. (D) (A-B) Proportion of proteins predicted correctly by AF2-M from a set of approximately 500 “unseen” proteins (A; no known structure, Foldseek TM score < 0.5 to any protein in the AlphaFold training set) and approximately 500 “seen-like” proteins (B; no known structure, Foldseek TM score > 0.5 to at least one protein in the AlphaFold training set), filtered by different pLDDT cutoffs for the predicted oligomeric complexes. For monomeric proteins, the pLDDT value used corresponds to the folded oligomeric state with the highest mean ipTM. Breakdowns are shown for the different folded oligomeric states as well as the overal total number of correct proteins. The number of correct proteins out of the total number of proteins in each bar is displayed in the graph. (C-D) Proportion of proteins predicted correctly by AF2-M from a set of approximately 500 “unseen” proteins (A; no known structure, Foldseek TM score < 0.5 to any protein in the AlphaFold training set) and approximately 500 “seen-like” proteins (B; no known structure, Foldseek TM score > 0.5 to at least one protein in the AlphaFold training set), filtered by different pLDDT cutoffs for the AFDB monomeric structure. Breakdowns are shown for the different folded oligomeric states as well as the overal total number of correct proteins. The number of correct proteins out of the total number of proteins in each bar is displayed in the graph. (E) Number of proteins per pLDDT cutoff per Foldseek TM-Score Bin for data shown in Fig. 4B.

**Figure S3.**
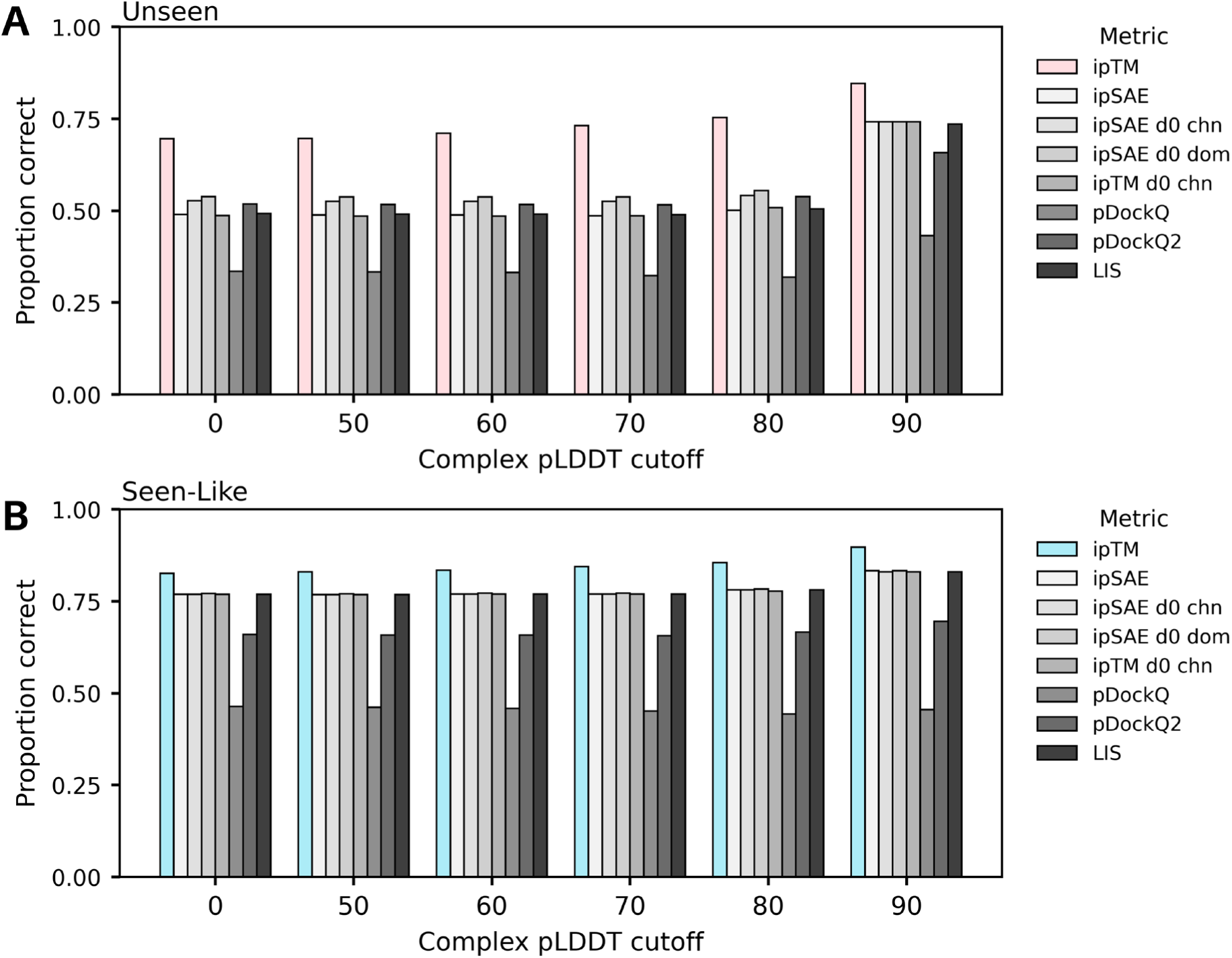
– Proportion of unseen and seen-like proteins predicted in the correct oligomeric state by AF2-M using different confidence metrics. (A-B) Proportion of proteins with (A) Foldseek TM scores < 0.5 (Unseen) or (B) > 0.5 to any protein (Seen-like) in the AlphaFold training set that were predicted correctly using different interface confidence metrics, including ipSAE, ipSAE d0 chn, ipSAE d0 dom, ipTM d0 chn, pDockQ, pDockQ2, LIS, and ipTM. Performance was assessed after filtering out structures based on pLDDT.

**Figure S4.**
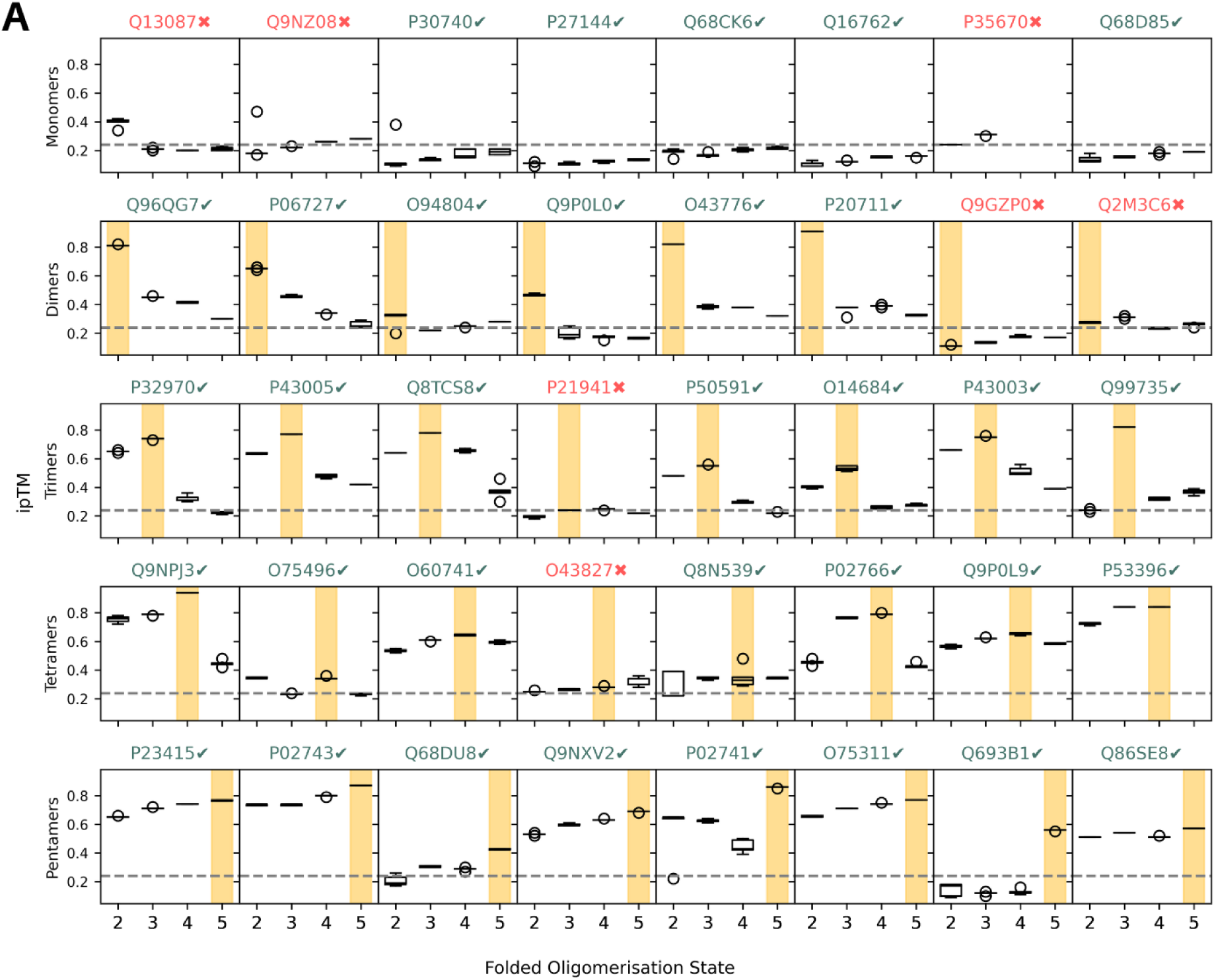
– ipTM distribution plots for AF3 predictions. (a) Box plots showing ipTM score distributions for each protein in different folded oligomeric states across the 40 test set proteins. The annotated oligomeric state highlighted in yellow. UniProt IDs are noted in green for correct predictions and red for incorrect ones.

**Figure S5.**
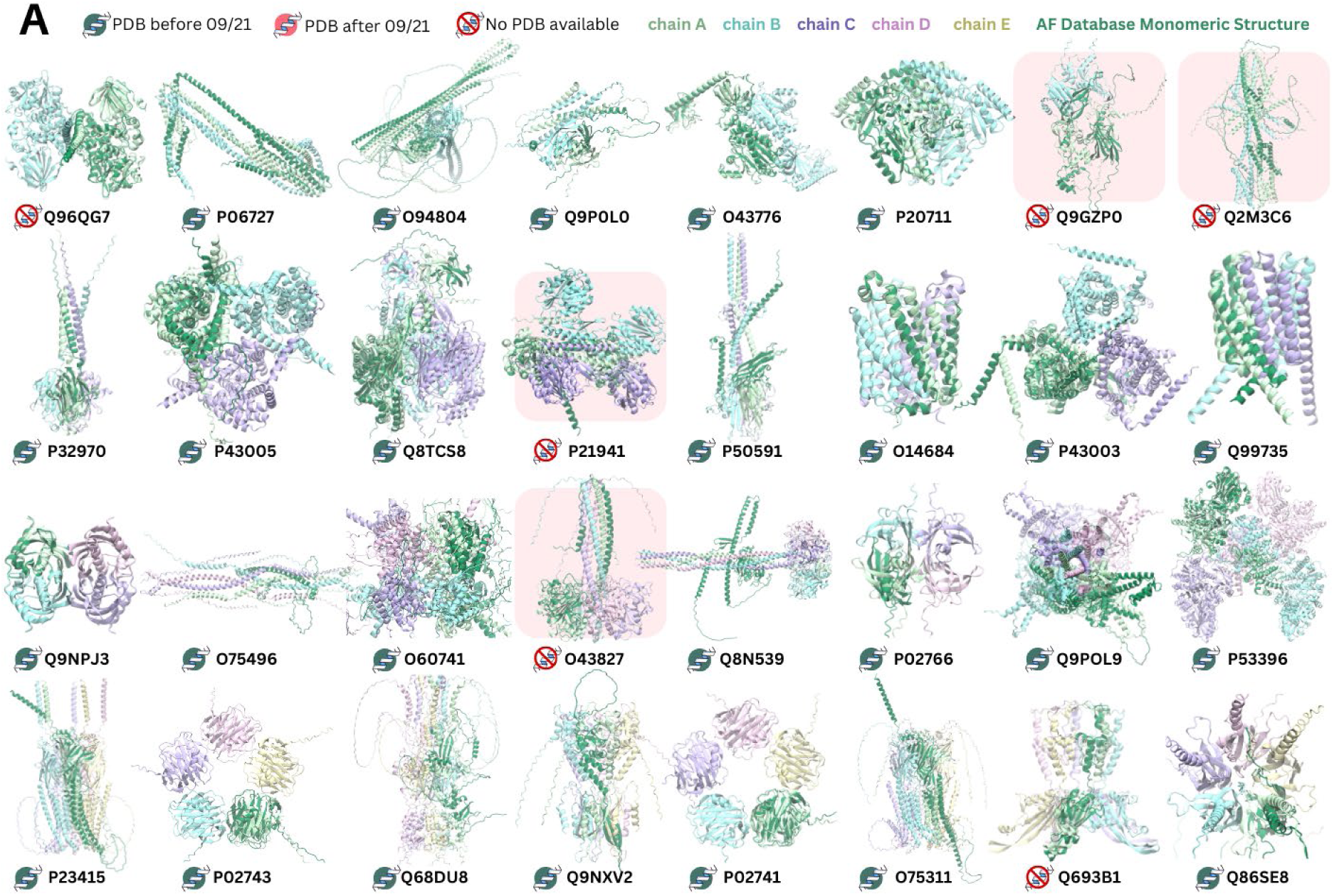
– structural images of AF3 predictions. (a) Snapshots of the top ranked structural predictions by AF3 in the correct oligomeric state for each multimeric protein (ribbon, coloured by chain), aligned with the AFDB prediction of the protein as a monomer (ribbon, dark green). The presence of an experimentally resolved structure in the PDB or AlphaFold3 training set is indicated with an icon near each UniProt ID. The structures of proteins predicted in an incorrect oligomeric state are highlighted in red.

